# Brain representations of visual short-term memories during healthy human aging: occipital decline and frontoparietal compensation

**DOI:** 10.64898/2026.10.26.761884

**Authors:** Daniel J. Mitchell, Cam-CAN, Rhodri Cusack

## Abstract

Visual short-term memory declines steeply with age, but the link to neural representations of memory content remains unknown. We used fMRI decoding to relate remembered motion-direction representations in visual and frontoparietal cortex to age and performance, in 115 adults, aged 25–87, from the Cambridge Centre for Ageing and Neuroscience (Cam-CAN) cohort. Decoding was strongest in visual cortex, but also significant in frontoparietal regions, with different profiles. Visual representations weakened with memory load and age, and statistically mediated age-related performance decline, especially during recall. This coupling, however, weakened with age: some older adults retained stronger visual representations than performance implied, suggesting impaired access to a partially preserved memory trace. Frontoparietal representations, instead, were largely insensitive to load and strengthened with age during maintenance, partially offsetting performance decline. Therefore, interventions to boost VSTM in older adults might target compensatory frontoparietal representations during memory maintenance, and readout of untapped visual representations at recall.

## Introduction

Our ability to hold information in visual short-term memory (VSTM) deteriorates markedly from early adulthood^1^. With visual short-term and working memory fundamental to much everyday cognition^2–4^, understanding this decline with age is critical, especially as lifespans increase^5^. Age-related declines manifest across multiple memory measures, including the quality, quantity, selectivity and feature-bindings of memory representations^6–10^. Nonetheless, older adults retain some attentional control over items in memory^11,12^, and continue to benefit from contextual cues at recall and metacognitive awareness of their memory performance^10^. An enduring question in memory science more broadly is how far forgetting reflects loss of the memory trace or impaired access to it^13,14^. In the context of aging VSTM, this remains unresolved, with direct investigation requiring neural measures of momentary memory content to be integrated with age-related differences in recall accuracy.

Over the past two decades, much has been learnt about VSTM representation in the human brain^15–20^. In younger adults, evidence points to roles of both frontoparietal cortex and early sensory cortex, but with vigorous debate surrounding their relative importance for memory representation^21–27^. In sensory cortex, memory contents are readily decodable from activity patterns, despite overall activation being low^28–33^. These early sensory representations relate to performance, with neural decoding accuracy declining along with recall precision as memory load increases^30,34^.

At the same time, frontoparietal activation increases with memory load^30,35,36^. Frontal activity may partially reflect actual or perceived difficulty, while parietal activity plateaus with memory load to track the limited number of stored items. Similar asymptotic responses in tasks without explicit memory requirements suggest that activation reflects attentional individuation of a limited number of items^36^. The role of frontoparietal regions in directly representing memory contents remains controversial, because while their activation patterns frequently distinguish memoranda^33,34,37–49^ decoding is often context dependent, weaker than in visual cortex, and sometimes undetectable^30,50^.

Thus, both sensory and frontoparietal regions jointly support VSTM representation, with somewhat complementary roles^22,46,47,51,52^. Critically, this balance may change with age^53^ and could therefore reflect age-related differences in behavioural performance. However, although numerous studies have examined neural responses during VSTM tasks in aging populations^11,53–61^, it remains unclear how declining VSTM performance with age relates to brain representation of memory contents. Canonical theories of frontoparietal involvement in aging are based largely on univariate over-recruitment, whose interpretation can be ambiguous without direct relation to memory content or behavioural performance^62–64^. Multivariate information has been proposed as stronger evidence for neural compensation^59,65^, but even multivariate studies have typically decoded general task variables (condition, load) rather than the memoranda themselves, and without relating these to behaviour^53,59^. Crucially, for neuroimaging data to provide practical insights for modifying age-related behavioural decline, the three-way relationship between age, cognition and brain measures is key to identifying potential mediation or compensation (suggesting mechanistic pathways that interventions might target) and moderation (situations where age-related differences in behaviour might be uncoupled from neural decline). If VSTM representations are maintained, but older adults lose access to them, we would expect to see neuro-behavioural uncoupling specifically at recall. In contrast, processes affecting the strength of memory representation should manifest earlier, during the maintenance period.

Here, we use fMRI and multivariate decoding to address this gap in the literature: we directly measure neural VSTM representations, across memory load and age, and relate these to behavioural decline over the healthy adult lifespan. Distinguishing two key brain systems (sensory, frontoparietal) and three task phases (encoding, maintenance, recall), we identify where and when neural representations are coupled to age-related behavioural variability in a manner potentially amenable to intervention.

## Results

To quantify neural representations of remembered stimuli, and relate these to individual differences in behavioural memory accuracy, we adopted a previously established fMRI paradigm using continuous delayed report of dot-motion directions. The stimuli and task (Figure 1A)were adapted from Emrich et al.^30^. On each trial, participants memorized the linear motion direction of 1-3 briefly presented dot patches, held these in mind for eight seconds, and then reported one direction specified by a colour cue. Participants were not told that on 90% of trials the probed patch moved in one of three critical directions.

**Figure 1.**
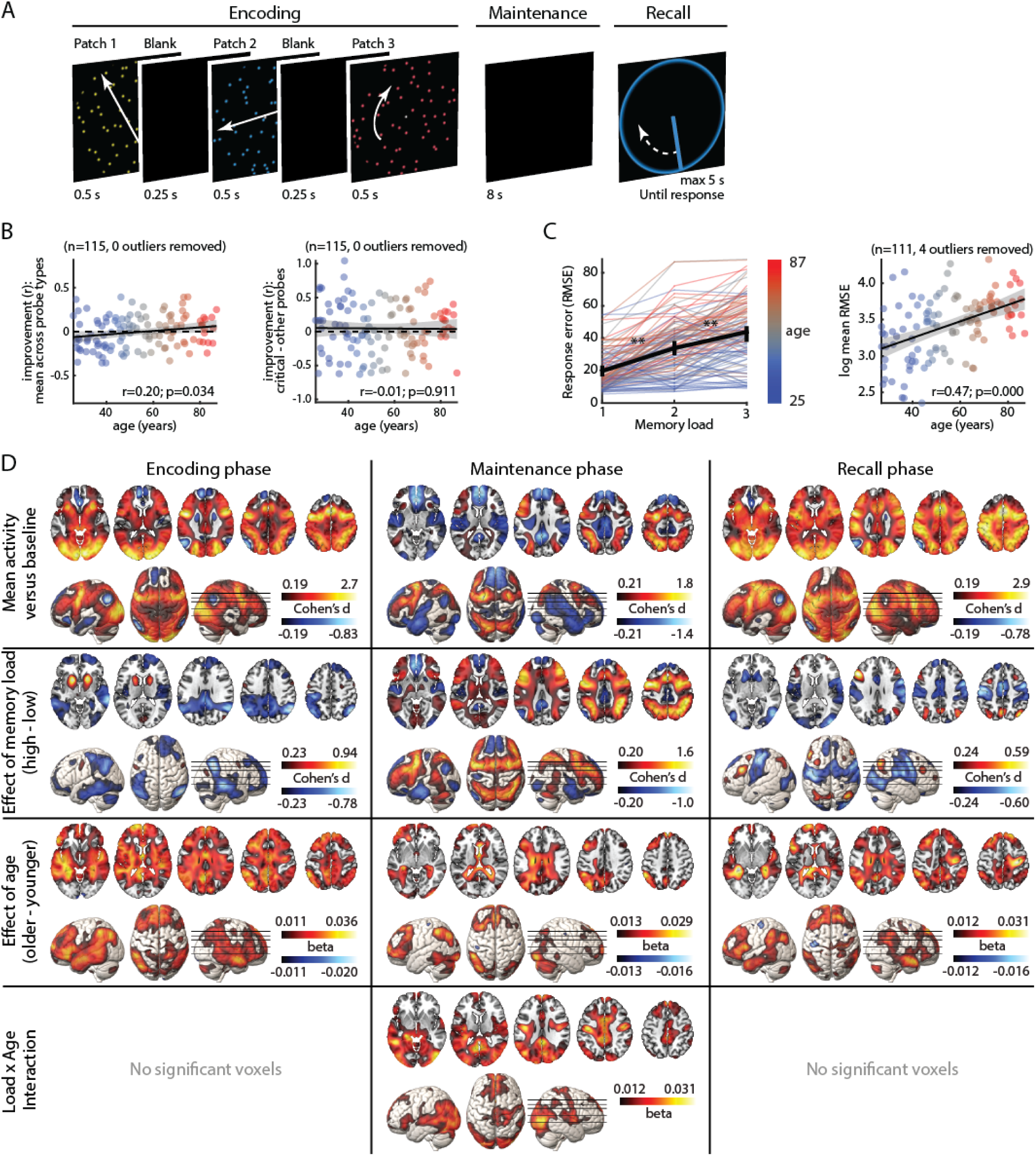
Task, behaviour, and univariate fMRI response. (A) Example of a single trial, with a memory load of 2. White arrows were not presented but are added here to indicate motion direction of the dot patches (solid arrows) and response pointer (dashed arrow). In this example, the translational motion of the yellow and blue stimuli is to be remembered, while the rotational motion of the red stimulus can be ignored. (B) Minimal improvement over trials, regardless of age, whether collapsing across probe types (left) or specific to critical-direction probes (right). (C) Increasing error with memory load (left) and age (right). In (B, C), error bars and grey bands indicate 95% confidence intervals. Asterisks indicate significant differences between adjacent conditions, **p<0.01. (D) Univariate fMRI effects of memory load and age. Each contrast is thresholded at FDR<0.025 per tail, overlaid on horizontal slices and surface renderings. Slice positions are marked on the right hemisphere rendering.

### Behaviour

We first tested for response bias as a function of the three critical probe directions (7, 127, 247 degrees, clockwise from 3 o’clock), collapsing across memory load. Responses to 7 and 127 degrees showed a significant clockwise bias, while responses to 247 degrees were biased anticlockwise (Extended Data Figure 1A). This is consistent with previously reported bias away from cardinal and towards oblique angles, in both perception and VSTM^66–68^. This “obliqueward bias”, averaged across the three directions, increased significantly with age (r=0.38, p<0.001) (Extended Data Figure 1B). These direction-specific response biases were subtracted from each response per critical direction before recalculating memory errors separately for each memory load.

To check that participants did not learn the three critical directions, for each participant we next correlated the squared error per trial with the trials’ position across the experiment, separately for trials with probes at critical and at other directions. A negative correlation thus indicates improvement over time, which we inverted so that positive values indicate improvement. A mixed effects linear model then tested whether improvement depended on the within-person factor of probe type (critical, other), the between-person covariate of age, or their interaction (Figure 1B). The main effect of age was weakly significant (F(1,113)=4.59, p=0.034), with older people tending to improve more over time; however, importantly, there was neither a main effect of probe type (F(1,113)=1.66, p=0.201) nor an interaction (F(1,113)=0.013, p=0.911), and therefore no evidence that participants learnt the critical directions.

As expected, overall root-mean-squared error (RMSE) increased with memory load (Figure 1C, left). Mixture modelling suggested contributions from lower memory precision, and lower probability of reporting the target direction (Extended Data Figure 1C). The decreased probability of target reports had contributions from both increased probability of uniform random guessing and increased probability of reporting an uncued (non-target) direction at loads 2 and 3.

To assess effects of age, response error was summarized by combining across memory loads to maximise sensitivity (as in ^10^), and then transformed to achieve a more Gaussian distribution. Overall RMSE increased with age (Figure 1C, right), along with deterioration of all mixture model parameters (Extended Data Figure 1D), mirroring findings on a test of colour VSTM completed by the same participants approximately 1.5 years previously^10^.

The similar effect of age across all mixture model parameters is consistent with the possibility that older adults make more errors of all types, but could alternatively reflect an inability of the mixture model to adequately decompose the error distribution from limited trials^69^. It is also consistent with the suggestion that all error types ultimately derive from a single underlying change in neural population activity^70,71^. Given this uncertainty, and the observation that the mixture model goodness-of-fit (log-likelihood) declines with age^58^, we took forward only the primary measure of overall error (RMSE) to relate individual differences in neural and behavioural responses.

### Effects of memory load and age on the univariate fMRI response

Whole-brain, voxel-wise univariate effects of task, memory load and age are rendered in Figure 1D. As expected^72^, compared to baseline, the task activated the frontoparietal multiple demand (MD) system^73^ throughout all task phases (Figure 1D, first row), along with visual cortex at encoding and recall, and somatomotor cortex especially at recall.

Effects of memory load (Figure 1D, second row) were most pronounced during memory maintenance, when activity increased with memory load throughout the frontoparietal MD system, as well as in visual cortex. Maintenance activity decreased with memory load in the default mode network (DMN)^74^. DMN activity similarly decreased with memory load at stimulus encoding, when increases with memory load were restricted to small foci in lateral frontal MD regions and visual cortex, alongside a prominent response in basal ganglia that may reflect attentional filtering^75^. We observed minimal load sensitivity of visual cortex during encoding, consistent with effective matching of motion energy across target and distractor dot patches, and with these early responses primarily reflecting visual input rather than memorized content. At recall, frontoparietal activity continued to increase with load, although less strongly than during memory maintenance. Somatomotor activity decreased with load, likely reflecting increased difficulty of response selection.

With increasing age (Figure 1D, third row), overall activity tended to increase across much of the brain and during all task phases, with minimal evidence for activity decreases. Areas of increased activity included parts of MD, DMN and visual cortex, but did not obviously correspond to the group-mean response patterns to task or load.

Finally, age strengthened the load effect during the maintenance period (Figure 1D, fourth row), although again this spanned parts of MD, DMN and visual cortex, without obviously corresponding to the group-mean effect of task or load. There was little evidence that the load effect decreased with age in any region, or that it changed with age at encoding or recall. This is potentially surprising, being in the opposite direction to prediction of the CRUNCH model of compensation^64^, although consistent with a recent report^60^.

### Effects of memory load and age on multi-voxel representation of memorised direction

The results thus far largely replicate known effects of age and memory load on behavioural performance and brain activation, establishing the validity and sensitivity of the paradigm, and setting the stage for the key analyses that test the effect of age on neural representations of visual memoranda and the three-way relationship with behaviour.

We next used multivariate pattern analysis, to decode the contents of VSTM from activation patterns within regions of interest (ROIs) in visual cortex and frontoparietal MD cortex. We classified the motion direction of the probed stimulus on the 90% of trials when this was in a critical direction (Figure 2). Considering first the memory maintenance period, consistent with previous reports^30,31^, we observed strong representation of remembered direction in early visual regions (Figure 2A, middle, purple bars), which decreased with increasing memory load (Figure 2B, middle, purple bars). Notably, however, representation of remembered direction was also significantly greater than chance in the intraparietal sulcus (IPS) and most frontal MD regions (Figure 2A, middle, green bars), albeit weaker than in visual ROIs. Interestingly, these MD representations appeared largely invariant to memory load (Figure 2B, middle, green bars).

**Figure 2.**
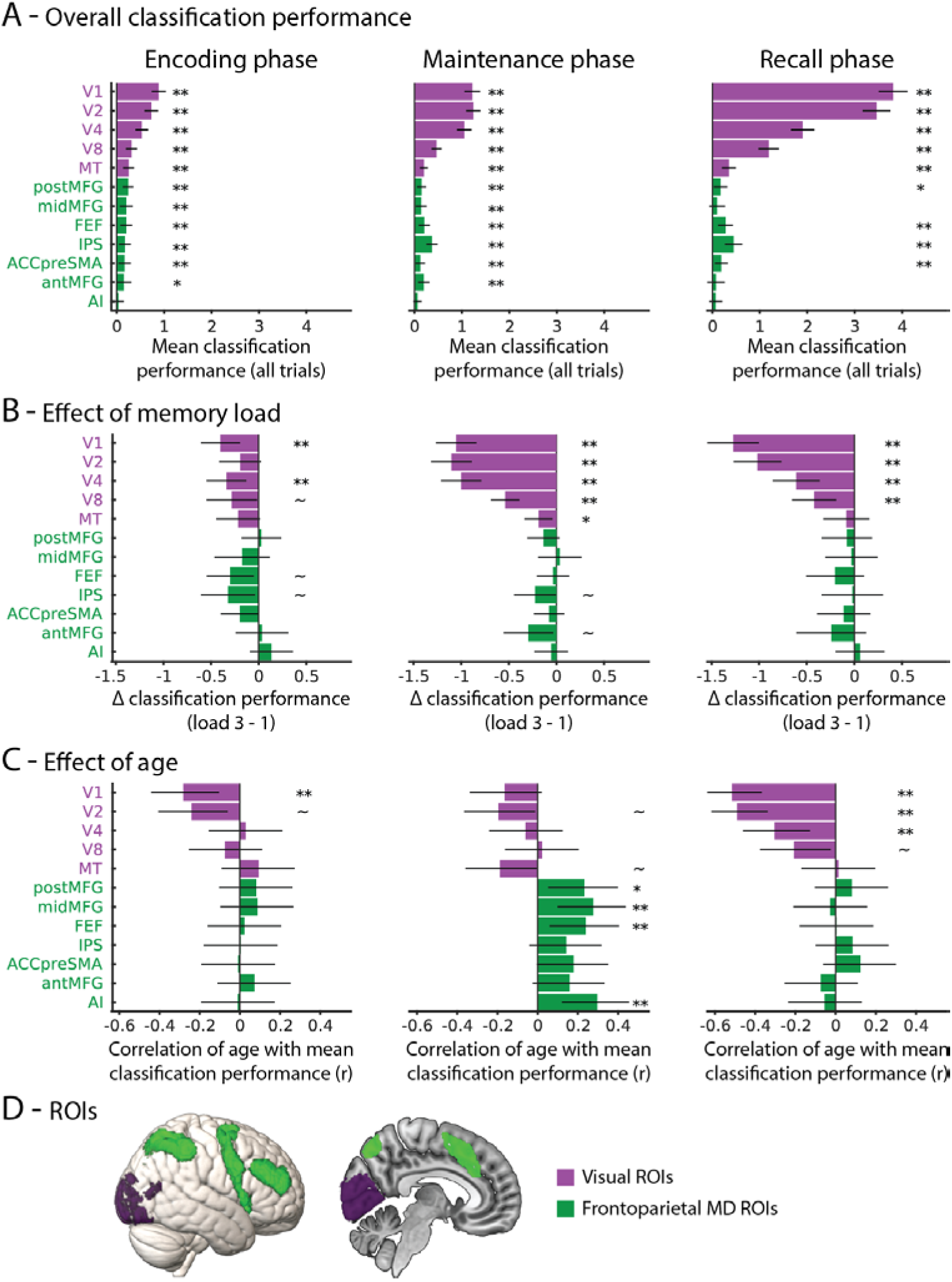
Multivariate decoding of memorized motion direction during each phase of the task. (A) Mean classification performance. (B) Effect of memory load. (C) Effect of age. Visual ROIs are highlighted in purple and frontoparietal MD ROIs in green. ** p<0.01 surviving FDR across ROIs; * p<0.05 surviving FDR across ROIs; ∼ p<0.05 but not surviving FDR across ROIs. Error bars show 95% confidence intervals. D) ROI locations, shown on lateral and medial views of the template brain.

During stimulus encoding, representation of direction showed a similar pattern (Figure 2A, left), strongest in early visual regions (purple bars), but also significant in all but one frontoparietal MD ROI (green bars). Visual representations weakened with increasing memory load during encoding (Figure 2B, left, purple bars), although to a lesser degree than during the maintenance phase, and only significant in some ROIs. Frontoparietal representations were again not significantly affected by load, after correction for multiple comparisons.

At recall, representation of remembered direction also showed a similar pattern (Figure 2A, B, right), again strongest in early visual regions, and also significant in the IPS, as well as several frontal MD ROIs. Once again, visual representations weakened with load, whereas MD representations did not.

Increasing age (Figure 2C) had a similar effect to increasing load at encoding and recall, but a strikingly different effect during the maintenance phase. Visual representations weakened with age, but most notably at recall, and non-significantly during memory maintenance. Frontoparietal MD representations instead appeared invariant to age at encoding and recall, but, interestingly, strengthened with age during the maintenance phase. No reliable interaction between age and memory load was observed in any region or task phase. Effects of age on visual cortex representations were consistent with a previous study by Carp et al.^53^, showing reduced distinctiveness at encoding and recall. Effects of age on frontoparietal representations differed from that study, showing a load-invariant strengthening of maintenance period representations rather than an interaction with load at encoding and maintenance. Alongside differences in task and precise anatomical correspondence, this difference may arise because the current analysis tests discrimination of visual motion direction across trials, whereas Carp *et al.* discriminated runs with visual stimuli from runs with verbal stimuli.

### Multi-voxel representation of other task features

Two supplementary decoding analyses addressed possible explanations for age-related weakening of memory content, beyond degraded integrity or accessibility of the to-be recalled representations themselves. Firstly, previous research suggests that attentional selection declines with age, such that distractors compete more strongly for limited short-term memory resources^7,8^. We therefore attempted to decode the distractor motion directions (clockwise versus anticlockwise). However, no reliable distractor representation was observed in any ROI at any task phase, neither was there any effect of age (all p > 0.05). Thus, we found no evidence in this dataset that weakened access to memory content in older adults depended on increased interference from irrelevant distracting information.

Secondly, we attempted to decode the colour of the probed patches. Since colour cued the to-be-reported direction, it should be perceptually represented at recall. This was the case in visual ROIs V1, V2, V4 and V8. During the encoding and maintenance phases, colour must be initially associated with direction, although it should be undecodable at memory load 3 when all colours are equally relevant and the identity of the probe is unknown. At load 1, colour representation also becomes unnecessary during the memory delay. However, colour could not be decoded from any ROI at any load prior to the recall phase. Colour representation did not reliably vary with age in any ROI at any task phase (all p > 0.05), suggesting that forgetting the to-be recalled directions at older age was unlikely due to forgetting the colours used to cue recall. Therefore, colour information was not analysed further.

### Relationships between neural representation, behavioural performance and age

Finally, and crucially, we assessed how neural representations during each task phase related to variability in behavioural performance, both within participants (i.e. correlation across memory loads, treating participants as a grouping variable) and between participants (treating memory load as a grouping variable), and then asked whether the neural representation mediated, compensated for, or interacted with the effect of age on individual differences in performance. We focused on the distinction between the visual and frontoparietal MD ROIs, averaging the response across individual ROIs within each set. The bivariate relationships between neural representation and behavioural errors are reported in Table 1, and illustrated in Figure 3A. Across both brain networks and all task phases, stronger neural representation was associated with less error in the memory response (consistently negative slopes). This relationship was significant in all cases, both across memory loads per person (individual lines), and across people per memory load, except for the MD representation at recall.

**Figure 3.**
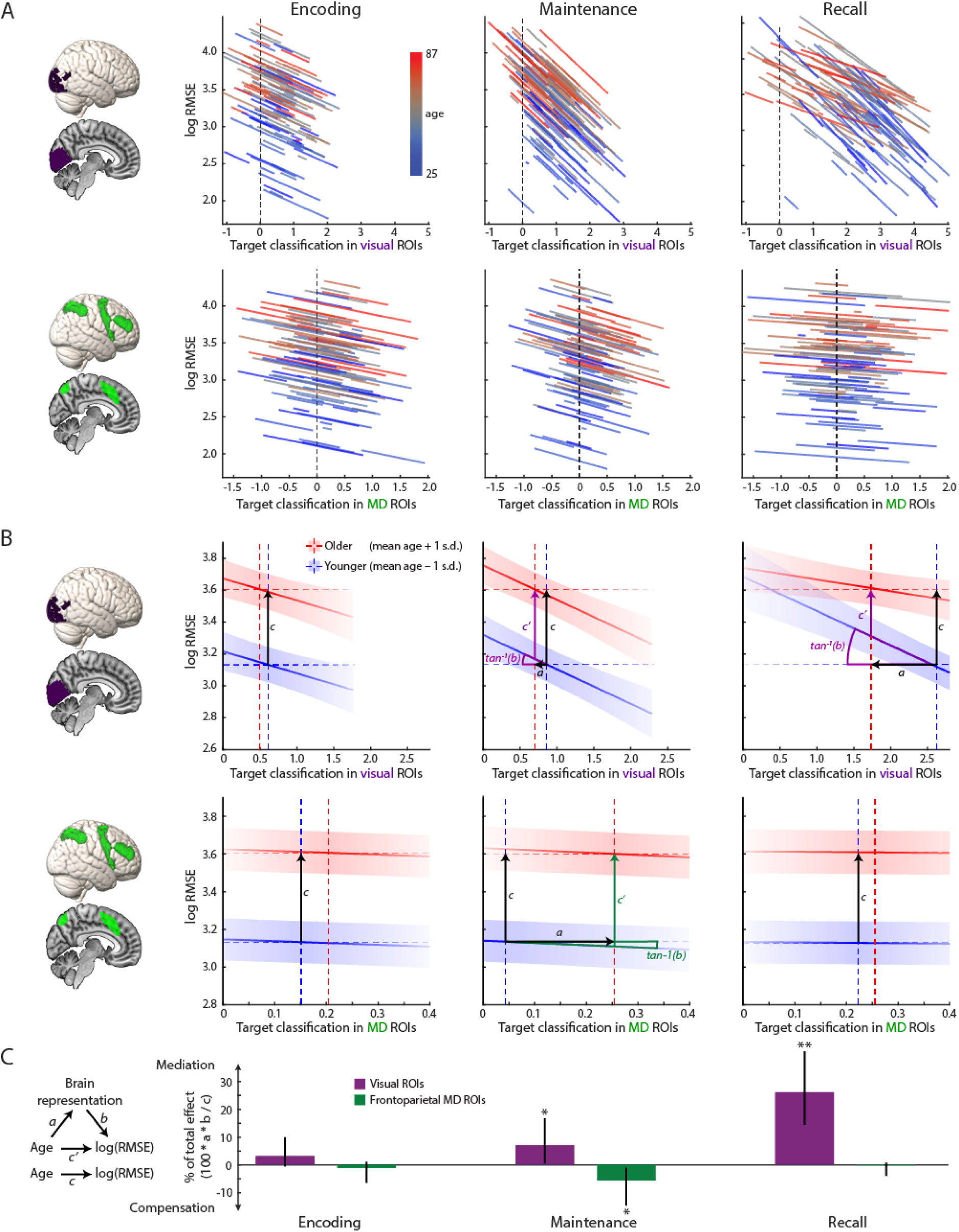
Relationships between multivariate representation of target direction, behavioural performance, and age. (A) For each network of interest (rows; ROIs rendered on left), and each task phase (columns), each plot illustrates the per-participant relationship between behavioural error and neural representation. Each line is plotted between the lowest and highest classification accuracy (across memory loads) for a single participant. Data are fit using parallel lines except for the one case (visual cortex classification during recall) when age interacted with the within-person slope (p<0.05), for which slopes are plotted as a function of age. (B) For each network of interest and each task phase, plots illustrate the modelled relationship between individual differences in behavioural error and neural representation, at younger and older ages (mean age +/- 1 standard deviation; blue and red respectively), at the mean memory load. Dashed horizontal lines, with separation labelled c, show the total effect of age on behaviour, irrespective of brain representation. Dashed vertical lines show the effect of age on the brain representation, irrespective of behaviour. Solid lines show the joint effect of age and brain representation on behaviour, excluding the interaction term unless significant (p<0.05). Error bands are 95% confidence intervals. (C) Bars quantify the percentage of the total effect of age on behavior that is statistically mediated, or compensated for, by the brain representation, assuming the causal model on the left (with memory load as a grouping variable, and the interaction term included if significant). Error bars are 95% bias-corrected bootstrap confidence intervals. ** p<0.01, * p<0.05. Where mediation or compensation are significant, vectors and angles corresponding to the a, b & c’ paths are plotted in panel B. In the case with an interaction, the indirect effect is conditional on age, and is only illustrated at the younger level.

**Table 1.**
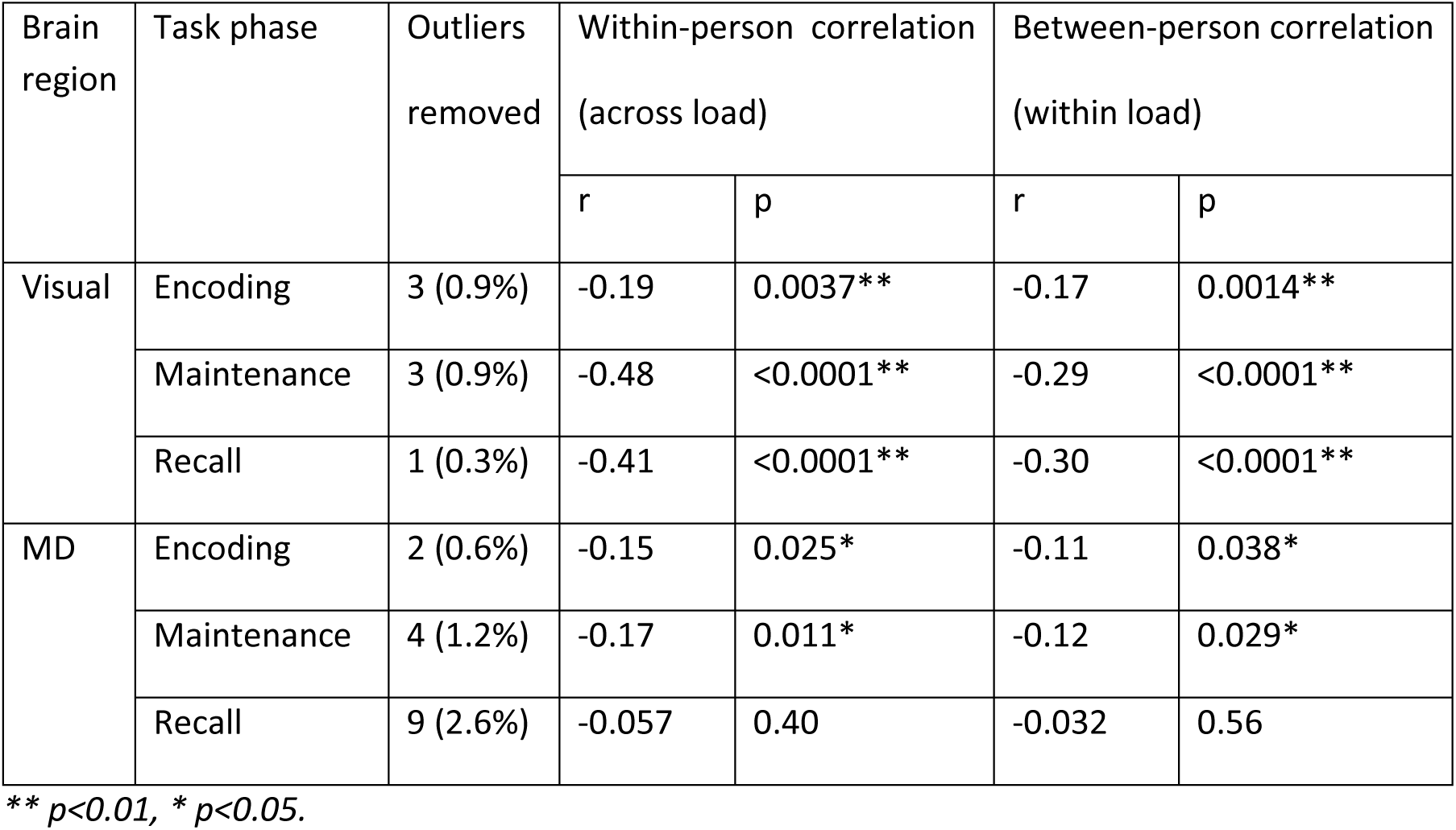
Bivariate relationships between neural representation and behavioural errors, for each brain region and each task phase.

The between-person correlations suggest that differences in the strength of neural representation could mediate, or compensate for, age-related differences in memory performance. To test this, we assessed the combined effect of (1) age on neural representation, and (2) neural representation on behaviour while controlling for age. The product of these coefficients can be interpreted as an indirect effect of age on behaviour that can be partly explained by the brain representation^76^. Joint effects of age and brain representation on between-person differences in behaviour are reported in Table 2 and illustrated in Figure 3B-C. Within visual cortex, the direction of the indirect effect matched the overall effect of age on behaviour, consistent with weaker representations partially mediating worse performance with age. This effect strengthened across the trial phases, being nonsignificant at encoding, weakly significant during the maintenance period, and strongly significant at recall. At recall, a significant interaction between age and visual representation also emerged, such that the mediation effect weakened with advancing age. This implies that as age becomes high enough, further declines in performance become progressively uncoupled from differences in visual representation. This raises the possibility that some older individuals may retain stronger representations in visual cortex than their poor performance would suggest.

**Table 2.** Joint effects of age and brain representation on between-person differences in behaviour.

| Brain region | Task phase | Outliers removed | Indirect effect of age on behaviour, via brain representation |  | Interaction of age and brain representation |  | Interpretation |
| --- | --- | --- | --- | --- | --- | --- | --- |
|  |  |  | % of total effect | p | F(1,330-8) | p |  |
| Visual | Encoding | 3 (0.9%) | 3.2 | 0.10 | 0.17 | 0.69 | Null |
|  | Maintenance | 3 (0.9%) | 7.1 | 0.032* | 0.74 | 0.39 | Mediation |
|  | Recall | 1 (0.3%) | 26.2 | 0.0030** | 9.84 | 0.0019** | Self-moderated Mediation |
| MD | Encoding | 2 (0.6%) | -1.1 | 0.29 | 0.086 | 0.77 | Null |
|  | Maintenance | 4 (1.2%) | -5.6 | 0.015* | 0.060 | 0.81 | Compensation |
|  | Recall | 9 (2.6%) | -0.2 | 0.47 | 0.24 | 0.63 | Null |

\*\* $p < 0.01$ , \* $p < 0.05$ .

A different pattern was observed in frontoparietal MD regions. While representational strength, adjusted for age, also correlated negatively with memory errors, in this case representation strengthened with age, to partially oppose the direct effect of age on performance. In other words, the total effect of age on performance was smaller than it would have been were it not for the greater neural representation in frontoparietal MD regions. This suppressor effect is consistent with functional compensation for the deleterious effect of increasing age.

## Discussion

In a large, population-based cohort, we characterise how neural representations of VSTM contents differ across the lifespan, and relate to differences in memory performance with age. With increasing age, representations in visual cortex were found to partly degrade (mediating behavioural decline), and partly become inaccessible (uncoupling from behaviour at recall), while frontoparietal representations were recruited during the maintenance period to compensate. Below, we highlight key insights and discuss potential implications.

### Complementary memory representations in visual and frontoparietal cortex have different susceptibility to ageing

Based on evidence from young adults, debate persists regarding the relative roles of visual versus frontoparietal cortex in representing VSTM content^21–23,27,77^. The present results support the consensus that distributed representations coexist across multiple regions, with complementary roles^17–19,46,51,52^. While we found memorised directions to be represented most strongly in visual regions, across all memory loads and task phases, weaker direction decoding was also significant in most frontoparietal MD regions. The differential strength of representation across ROIs is difficult to interpret: frontoparietal representations may appear weaker due to finer-scale intermixing of neurons, which are less visible to fMRI than topographic organizations in visual regions^78^.

During memory maintenance, these visual and frontoparietal representations responded differently to within-participant changes in memory load and between-participant variation in age. Visual representations weakened with load, matching previous reports^30,34^, whereas frontoparietal representations appeared largely load-invariant. Importantly, while visual representations weakened with age primarily at recall, frontoparietal representations appeared age-invariant at recall, and strengthened with age during memory maintenance. Thus, while age-related neural dedifferentiation^79^ was substantial in visual regions at recall, it was not ubiquitous, either across brain networks or task phases. Overall, these observations are consistent with visual representations being relatively fragile, and frontoparietal representations more robust^41^. The differential age effects across brain networks also resonate with the behavioural finding that older adults’ diminished VSTM capacity primarily affects domain-specific sensory stores, alongside preserved domain-general capacity^80^.

### Representational decay in visual cortex partially mediates age-related behavioural decline, yet residual representations may hold latent potential in older adults

Supporting previous findings, the strength of memory representation in visual cortex correlated with memory performance, both within participants^30,31^, and across participants^37,81^. We identified two, noteworthy, age-related properties of this relationship. Firstly, the behavioural variance explained by the neural representation increased throughout a trial, being greater during maintenance and recall than during encoding, and with neural mediation of the age effect increasing from 3% at encoding, to 7% during maintenance, to 26% at recall. Sometimes, perhaps especially when memory capacity is exceeded, individual differences at encoding can constrain performance^72,82,83^, consistent with capacity limits manifesting at an early attentional selection stage^36,84^, regardless of age^85^. The present data highlight that the integrity of visual representations at memory recall may also be critical to performance, consistent with either accumulating representational instability^86^ or interference from the probe display^87^, especially with increasing age.

The second key observation from visual cortex is that the slope of the relationship between behaviour and occipital representation at recall depended on age, with performance strongly coupled to visual representation in youth but less so at older ages. This suggests that visual cortices of some older individuals might preserve reliable VSTM representations, which, for some reason, become less effective in driving accurate performance than they would have been in younger individuals. This could reflect a behavioural bottleneck downstream of visual representation. One possibility is that older individuals rely more heavily on prior expectations, which they allow to override the available sensory evidence^88^. This resonates with the increased obliqueward bias observed in older adults. Despite being detrimental in this task, such a strategy could be adaptive in more generally, if older people compensate for noisier inputs by applying a lifetime of experience to make plausible predictions. A related possibility is that older adults are less confident in the accuracy of their internal representation^10^, leading to a lower threshold for guessing. Either way, an appealing interpretation is that older participants may retain untapped neural potential, raising the possibility of improving their VSTM performance if one could re-establish more efficient readout of residual memory representations in visual cortex.

A potential interpretational concern is that the neural pattern at recall partly reflects perceptual representation of the response pointer, alongside memory representations that drive the response. However, the form of the interaction with age in predicting performance argues against this perceptual explanation being critical for the age effects: older participants tended to perform worse regardless of the strength of their visual representation at recall (Figure 3B, top right); therefore, reduced accuracy in pointer position was not necessarily reflected in weaker visual representation. Here, the stimulus encoding period can also be considered as a control condition, when there was no evidence that visual representations, presumably also perceptually dominated, mediated age-related effects on memory performance (Figure 3B, top left), Together, these observations suggest that the age-related vulnerability is likely to lie primarily at the level of maintaining and accessing memory representations, rather than perceptual fidelity.

### Compensatory frontoparietal representations during VSTM maintenance buffer age-related behavioural decline

Unlike visual regions, where memory representation and its mediation of age-related memory errors strengthened from encoding to maintenance to recall, performance was only associated with frontoparietal MD representation during encoding and maintenance. This again suggests differences in the roles of frontoparietal and visual representations: while both may support memory maintenance, frontoparietal regions are linked to performance at encoding more than readout, whereas the converse is observed for visual regions.

During memory maintenance, we observed “inconsistent mediation” of age-related behavioural decline by strengthened frontoparietal MD representation. This suggests that frontoparietal representation acts as a suppressor variable, with a compensatory effect that helps to offset the direct deleterious effect of age on performance. An often-reported finding is a posterior-to-anterior shift in neural activation with increasing age, which might reflect compensatory processing or reduced neural efficiency^62^. This pattern was not obvious in our univariate data, where activation tended to increase with age across both anterior and posterior brain regions. A previous report has also suggested that multivariate prefrontal activity in older age does not become more informative than visual activity in coding memory load^59^. The current findings suggest, however, that frontoparietal MD regions can play an age-related compensatory role in maintaining a representation of the target feature itself. Thus, these results are potentially consistent with a compensatory “posterior to anterior *representational* shift in aging,” although we would characterise it more as a sensory to frontoparietal shift. Previous research in young adults indicates that perceptually challenging situations recruit MD representations when visual representations are weakest^89^. Current results suggest that enhanced MD representation might similarly compensate for reduced visual fidelity across individuals^90^, and, importantly, might buffer short-term memory errors with advancing age.

These compensatory frontoparietal representations may store information in a transformed or abstracted format^91^, such as verbal labels^92,93^, reflecting a shift away from a visual imagery strategy^94^. Speculatively, this could explain the restriction of the effect to the maintenance period, since a native visual format dominates at encoding^95^ and is encouraged at recall by the nature of the final response. Several recent findings suggest that manipulations associated with better short-memory remain effective into old age, despite overall memory decline^10–12,85,96^. Augmenting VSTM via auxiliary MD representations may represent a neural manifestation of such a spared strategy.

It is interesting that frontoparietal MD activity can partially compensate for age-related decline in VSTM despite partially mediating age-related decline in fluid intelligence^97^. Perhaps any MD dysfunction is deleterious in fluid intelligence tasks for which MD regions are essential, whereas MD engagement may be just one of multiple strategies for successful VSTM performance^57,94^, so recruitment of an underperforming system can still be beneficial.

### Limitations and open questions

Within-subject weakening of remembered direction decoding at higher memory loads could reflect greater classifier difficulty in distinguishing multiplexed patterns, as well as weaker representations *per se*. While these alternatives are hard to disentangle (see ^30^ for further discussion), this issue does not bear on the key results concerning effects of age.

We were unable to decode the colour of memory items prior to probe, or the rotation direction of distractor items, although the task requires some representation of colour, and susceptibility to distraction is believed to increase with age^7,8^. The lack of robust colour decoding may reflect the relative simplicity of the colour cue in this task, comprising only three easily verbalised values. Distractor representation may have been undetectable because the rotating distractors were consistently irrelevant and thus easily ignored, unlike studies where distractors resemble memoranda and change relevancy trial-by-trial. Even then, the human brain appears remarkably successful at suppressing task-irrelevant information^28,42,98^. Age-related susceptibility to distractors may also be minimal at low memory loads^7,99^, transient and thus hard to detect at the temporal resolution of fMRI^8,100–102^, or may impact memory performance without requiring storage of the distractors themselves^103^. Investigating age effects on neural representations of cue and distractor information may require designs in which cues are harder to represent and distractors are harder to suppress.

Although the experimental design allowed some separation of task phases, it was not optimised for this purpose. Interesting differences between phases were observed, suggesting some success in separating memory encoding, maintenance and recall processes. However, future experiments could usefully replicate the current findings in designs with longer or more variable inter-phase intervals, and that explicitly separate recall from response.

Here, we characterised age effects across memory loads that are generally within capacity. However, people may engage different strategies as memory capacity is exceeded and performance breaks down^72,85^. These factors correlate with fluid intelligence^104^, which declines with age^97,105^, and although older adults can use a selective strategy when instructed, they struggle to judge its effectiveness^85^. Therefore, it is uncertain how far the current findings generalise to memory loads that substantially exceed capacity.

Finally, the current cross-sectional data conflate within-individual aging processes with potential generational differences across age. The finding that between-individual covariance between neural representation and behaviour (Figure 3B) mirrored within-individual (across memory load) covariance (Figure 3A) suggests that within-person and between-person effects might reflect a common mechanism. Nonetheless, any specific interpretation in terms of aging mechanisms would require replication with longitudinal data.

### Conclusions

The current results advance understanding of how, where and when VSTM representations in the human brain change across the adult lifespan, and how these changes relate to declining memory accuracy with age. Results suggest that a major locus of performance failure in older adults may arise at memory recall, partly mediated by fragility of visual cortex representations, but with additional contributors downstream. The findings are encouraging in suggesting that older adults recruit compensatory frontoparietal representations during maintenance to buffer performance decline, and that older adults might retain more intact visual representations of remembered items than their behaviour would indicate, raising the possibility that performance might be improved if effective readout of these representations could be re-established. Focusing on neurocognitive mechanisms that operate at these stages may be fruitful in the quest to ameliorate performance decline in healthy old age.

## Methods

### Participants

Participants reported here are a subset from the population-based healthy adult cohort recruited at Stage 3 of the Cambridge Centre for Ageing and Neuroscience project (see ^106^ for full details of the sample and exclusion criteria, and ^107^ for a recent review). All participants performed various psychological and neuroimaging assessments, and we focus here on analysis of the fMRI VSTM task. Data from this task have been reported previously, to address different questions^57–59^. 125 participants performed the fMRI VSTM task. Two of these were excluded due to incomplete EPI data; a further eight were excluded due to poor performance when memorizing a single item (mean absolute error >30 degrees). The final sample thus consisted of 115 participants (54 female) with ages ranging from 25 to 87 at the time of fMRI (mean 54.5 years; approximately equal numbers per decile). The study was conducted in accordance with ethical approval from the Cambridgeshire 2 (now East of England—Cambridge Central) Research Ethics Committee (reference: 10/H0308/50), and participants gave written, informed consent.

### Stimuli and task

Stimuli were presented on a 1920 x 1080 pixel screen, using Matlab (The Mathworks Inc.) and Psychtoolbox-3^108^. The screen had a width of approximately 20 degrees of visual angle (dva) when viewed through a mirror mounted on the head coil. Stimuli were displayed on a black background. To minimise eye-movements, participants were asked to fixate a central grey dot (diameter 0.22 dva) throughout the experiment. To minimise carry-over from previous memory representations, each trial was preceded by a 7 s blank inter-trial-interval (ITI). For the last two seconds of the ITI, the fixation dot turned from grey to white, to alert the participants that the stimuli were about to appear.

To manipulate memory load while equating visual stimulation, on each trial three coherently moving dot patches were then presented sequentially, one red, one yellow and one blue, in random order. Motion direction of one patch was to be reported after a brief delay, with a coloured post-cue serving to identify which direction to report. Colours were chosen to be easily discriminable, including by any participants with red-green colour-blindness. Each dot patch was viewed through a circular aperture (diameter 11 dva) and was composed of dots of diameter 0.26 dva, positioned pseudo-randomly at a density of 0.7 dots per dva^2^. Dot patches were presented for 500 ms, separated by 250 ms blank periods. To manipulate memory load, between one and three of the dot patches moved linearly at a speed of 2.75 dva/s; participants were asked to memorise these motion directions and the associated patch colours. To approximately equate visual stimulation across memory loads, the remaining patches rotated around the fixation point, at the same average speed, and were to be ignored. Thus, the memory load varied from one to three directions. For the purpose of decoding distractor representation, the rotation (clockwise, anticlockwise) of any distractor patches was fixed within a trial, and alternated across trials of a given memory load.

Following the disappearance of the third dot patch, participants were to keep the memorized motion directions in mind over an 8 s blank interval. This duration has previously been shown sufficient to partially dissociate neural representations of the memoranda from representation of the stimuli themselves^30^. A probe display then appeared, composed of a circle (diameter 11 dva) and a line connecting the fixation point to a random point on the circle. To cue which memory to report, the colour of the circle matched the colour of one of the memorized dot patches, and participants were instructed to report the motion direction of the corresponding dot patch by rotating the line to point in this direction. To provide their response, participants used a button box held in their right hand. Holding down either their index or middle fingers caused the pointer to rotate either clockwise or anticlockwise, with the mapping randomised across trials. When participants were satisfied that the line pointed in the correct direction, they pressed a third button with their right little finger. As soon as the participant pressed the third button, or after 5 s, the probe display disappeared, and the next ITI began.

To allow decoding to be cross-validated across independent data partitions, trials were presented across three runs, each run consisting of 30 trials. To combine data across repeated presentations of a given motion direction, on 27 trials per run, the probed dot patch moved in one of three critical directions (7, 127, or 247 degrees, clockwise from three o’clock). Participants were not told of this in advance. All other dot patches moved in random directions, including memorized but unprobed patches on critical trials, and all memorized patches on the remaining three trials. Across the 27 critical trials per run, memory load (one, two, or three directions) was counterbalanced with the probed patch’s motion direction (7, 127, or 247 degrees) and sequential position (first, second, or third). The colour of the probed patch (red, yellow, or blue) was also balanced and crossed with sequential position using a Greco-Latin square. The remaining three trials of each run included one of each memory load, with other parameters chosen at random. All trials within each run were presented in random order.

To familiarise participants with the task, before entering the scanner, participants practised five trials with no time limit, followed by five trials with a 5 s time-limit. This was repeated if necessary until the participant and experimenter were happy that the task was understood and responses could be given within 5 s. Participants were asked to report their best guess when unsure of the answer. Inside the scanner, participants received a couple more practice trials before starting data collection.

### fMRI acquisition

To measure event-related brain responses, during each of three functional runs while the participant was performing the task, T2*-weighted Echo Planar Images were acquired using the following parameters: Repetition time (TR) = 2 s, three echo times (TE = 12 ms, 25 ms and 38 ms), flip angle = 78 degrees, 34 axial slices of 2.9 mm thickness acquired in descending order with an inter-slice gap of 20%, in plane field of view (FOV) = 224 mm × 224 mm, and voxel-size = 3.5 mm × 3.5 mm × 3.48 mm. There were a variable number of scans per run, depending on reaction time, ranging from 294 to 349 (median 320).

To enable normalisation of individual brains to template space, a structural image was acquired for each participant, on a different day (median 49 days apart, range 75 to 1039). This used a T1-weighted 3D Magnetization Prepared Rapid Gradient Echo (MPRAGE) sequence with the following parameters: TR = 2250ms, TE = 2.98 ms, inversion time = 900 ms, flip angle 9 degrees, FOV = 256 x 240 x 192 mm, voxel size = 1 mm isotropic, and GRAPPA acceleration factor 2.

### Behavioural analysis

The task design allowed memory accuracy to be measured on a continuous scale per trial, with response error scored as the angular difference between the true and reported motion directions. This was summarized across trials as the root-mean-square error (RMSE). In principle, short-term memory can fail in several ways: a memory may become less precise, be lost entirely (leading to a guess), or be attributed to the wrong item (mis-binding). Mixture models of the response-error distribution aim to separate these sources of error, but they make strong assumptions and can be unreliable with limited trials^69^. By encapsulating all sources of error, with minimal assumptions, RMSE is expected to be more robust than individual mixture model parameters. We therefore used RMSE as our primary performance index, and used the mixture model mainly to correct for systematic response bias, as described below.

Mixture models fit to the error distributions, provided estimates of memory bias (average offset from probed direction), quality (kappa, the precision of each direction held in memory), quantity (K, the expected number of directions stored) and mis-binding (the likelihood of reporting a memorised direction but from an unprobed dot patch). We used a mixture model as introduced by Zhang and Luck^109^, and augmented by Bays, et al. ^110^, which included: a uniformly distributed component to model guesses, a von-Mises distributed component to model responses centred on the probed direction and further von-Mises distributions centred at the motion directions of unprobed dot patches, to model mis-binding responses. The location parameter of the von-Mises distribution (μ) was allowed to vary for the probed direction, but was fixed at zero for any unprobed directions, to equate the number of estimated parameters across memory loads 2 and 3. Model parameters were estimated using code adapted from Bays, et al. ^110^, as in Mitchell et al.^10^.

The mixture model was first fit per direction per participant (combining memory loads), for the three critical directions that were probed on 90% of trials. For this initial fit, we focused on the μ parameter, which we expected to deviate consistently from zero, indicating a response bias away from cardinal (horizontal and vertical) directions. Such an obliqueward bias had been demonstrated in perceptual^66^ and VSTM tasks^67,68^, and adjusting for this non-uniformity across the stimulus space can improve the model fit^68^. In a second step, RMSE and the mixture model were therefore recalculated per memory load per participant (combining directions), after adjusting each response for the estimated bias per direction. The bias-corrected RMSE from this second step served as the primary behavioural measure for subsequent analyses. Mixture model component parameters are reported for completeness in Extended Data Figure 1.

For each memory measure, performance was summarised across memory loads, using the mean across loads for RMSE and precision, the minimum across loads for the probability of target responses, and the maximum across loads for the probability of guesses and misbinding errors, to reduce the contribution of ceiling and floor effects for the mixture weights at lower loads as in ^10^.

### fMRI preprocessing

Analyses used SPM12 (http://www.fil.ion.ucl.ac.uk/spm), Matlab (The Mathworks Inc.), and “automatic analysis” software^111^. The structural images were bias-corrected, segmented and warped to match the MNI152 (Montreal Neurological Institute) template brain using SPM. The first five functional scans from each run were discarded to allow the signal to reach equilibrium. The remainder were combined across echoes at each TR (mean, weighted by estimated T2* contrast), spatially realigned to each other to adjust for head motion, interpolated in time to correct for the different slice acquisition times, rigidly coregistered to the structural image, transformed to MNI space (using the transform from the structural image), and resliced to 2 x 2 x 2 mm resolution. For voxel-wise univariate analyses, data were spatially smoothed using a Gaussian kernel of 10 mm full-width-half-maximum; for all other analyses, data were left unsmoothed.

### Regions of interest

To test the hypothesis that memory representations in early visual and frontoparietal cortex might be differentially vulnerable to aging, we selected regions of interest (ROIs) in each of these two networks. In visual cortex, we used the multimodal parcellation from the Human Connectome Project^112^ to define V1, V2 and MT ROIs, where Emrich, et al. ^30^ had observed strong representation of remembered direction. We also added V4 and V8 (also known as VO1) from the same parcellation, due to their known sensitivity to colour^113^, which we also aimed to decode. ROIs were generated from https://identifiers.org/neurovault.image:30759, dilated to fill a grey-matter mask.

In frontoparietal cortex, we defined ROIs based on the multiple-demand (MD) system^73^, whose response increases with cognitive demand across many tasks, and appears critical to fluid intelligence differences across age^97^. ROIs were defined based on data from Fedorenko et al.^114^; their figure 2, selecting frontoparietal regions responsive to cognitive demands across seven diverse tasks, divided into contiguous regions as described in Mitchell et al.^115^. These MD regions include the intraparietal sulcus (IPS), anterior insula (AI), anterior cingulate cortex/pre-supplementary motor area (ACCpreSMA), frontal eye fields (FEF), and three foci along the middle frontal gyrus (MFG); they are available at http://imaging.mrc-cbu.cam.ac.uk/imaging/MDsystem.

### fMRI modelling

The signal time-course, per voxel, per participant, was fit using two models. Each model distinguished three phases of each trial: the stimulus “encoding” phase, defined as the 2 s from the onset of the first stimulus to the offset of the third stimulus; the memory “maintenance” phase, defined as the 8 s delay from the offset of the third stimulus to the onset of the probe; and the “recall” phase, defined as the period from probe onset until response (or the disappearance of the probe if no response was made within 5 s). A first model, for univariate analyses, used three regressors for each phase, separating trials only by memory load (1, 2 or 3 directions). A second, more comprehensive model, for multivariate analyses, used separate regressors for each phase of every trial separately. All regressors were convolved with the canonical haemodynamic response function. Both models also included additional unconvolved regressors to model the six estimated motion parameters and the run means. Both models, and the data, were temporally high-pass filtered using a cut-off of 128 s.

### Multivariate pattern analyses

Multivariate pattern analyses used The Decoding Toolbox^116^ to classify features of interest per trial per participant, based on local patterns of coefficient estimates from the model described above. Following Emrich, et al. ^30^, we focused on the motion direction of the probed stimulus on the 90% of trials when this was in a critical direction (7, 127, 247 degrees). Results were averaged across three binary classifiers that distinguished each possible pair of directions. We also attempted to classify the motion direction of the to-be-ignored distractor stimuli (clockwise, anticlockwise) at memory loads 1 and 2, and the colour (red, yellow, blue) of the probed stimulus. For consistency with Emrich, et al. ^30^, we used an L2-regularised logistic regression classifier, as implemented in the LIBLINEAR package^117^. Also matching Emrich, et al. ^30^, classifiers were trained across all memory loads but tested separately on left-out patterns at each load. Classifier accuracy was assessed using leave-one-run-out cross-validation, i.e. each classifier was trained using patterns from two runs and tested on patterns from the third run, averaging results across the three folds. As the outcome measure, we used the signed distance to the discriminating hyperplane, which is randomly distributed around zero under the null hypothesis of no difference between classes; as an estimated distance metric, similar to the linear discriminant contrast, this has the advantages of being continuous, unbounded, and insensitive to differences in pattern variance^118,119^.

### Relating age, brain and cognition

Representation of target direction, separately averaged across visual cortex and frontoparietal MD ROIs, was related to behavioural performance (RMSE) using general linear models that isolated within-person effects driven by memory load (as in^30,31^), using the method of Bland and Altman ^120,121^. We analogously isolated between-person effects by treating load as the grouping variable.

Finally, we tested whether the data were consistent with the neural representation mediating, or compensating for, the between-person effect of age on performance^76^. To do this, we tested for joint significance of both an effect of age on neural representation and an effect of neural representation on behaviour while controlling for the main effect of age^122^. As above, load was modelled as a grouping variable. The product of these coefficients quantifies the effect of age on performance that can be attributed to an indirect causal path via the brain response. This indirect effect was expressed as a percentage of the total effect of age on performance (when the brain response is not included in the model). If this percentage is positive, the indirect path explains (some of) the total effect of age, consistent with the neural response (partially) mediating the age-related decline in performance. If the percentage is negative, the indirect path opposes the direct effect of age (sometimes termed inconsistent mediation, or suppression)^123^, consistent with the neural response compensating for age-related decline. By adding the interaction between age and neural representation when predicting behaviour, we also tested whether age moderated the between-person relationship between brain and behaviour.

### Statistical analyses

All analyses were performed in Matlab (The Mathworks Inc.). Across the analyses described above, we used paired two-tailed t-tests to compare conditions, one-tailed, one-sample t-tests to test for above-chance classification, Pearson correlation to test for linear associations, and general linear models to test joint effects of multiple variables.

Univariate fMRI analyses tested for an effect of memory load by contrasting coefficients for high and low memory load (3 vs 1 directions), and an overall effect of task by contrasting the mean of all memory loads versus the implicit fixation baseline. At the group level, these contrasts were tested against zero using two-tailed one-sample t-tests, and regressed against age to assess the main effect of age and age-by-load interaction.

For both univariate and multivariate analyses, multiple comparisons across voxels or ROIs were accounted for by controlling the expected false discovery rate (FDR)^124^ to be less than 0.05. Brain renderings used MRIcroGL software^125^.

When testing associations with behaviour, the behavioural variables were transformed to achieve approximate normality. Specifically, RMSE and kappa are non-negative and positively skewed and therefore a log transform was applied; mixing components for the target, non-target and guessing probabilities are bounded between zero and one, so an arcsine square-root transform was applied. Outliers were excluded per analysis based on a threshold of three scaled median absolute deviations from the median, as implemented by Matlab’s “isoutlier” function. Significance of mediation/compensation, and associated confidence intervals, were estimated via bias-corrected bootstrap, with 1000 samples.

## Acknowledgements

The Cambridge Centre for Ageing and Neuroscience (Cam-CAN) research was originally supported (2010-2015) by the Biotechnology and Biological Sciences Research Council (grant number BB/H008217/1); it was subsequently supported by the Medical Research Council (MRC) Unit programme grant MC_UU_00030/8, with a contribution (2017-2022) from the European Union Horizon 2020 Research and Innovation Programme (LifeBrain) Grant Agreement 732592. Daniel Mitchell was supported by the MRC Unit programme grant MC_UU_00030/7. We are grateful to the Cam-CAN respondents and their primary care teams in Cambridge for their participation in this study.

The Cam-CAN corporate author consists of the Project principal personnel: Richard N Henson, Lorraine K Tyler, Kamen A Tsvetanov, Carol Brayne, Edward T Bullmore, Andrew C Calder, Rhodri Cusack, Tim Dalgleish, John Duncan, Fiona E Matthews, William D Marslen-Wilson, James B Rowe, Meredith A Shafto, Marta Correia; Research Associates: Karen Campbell, Teresa Cheung, Simon Davis, Linda Geerligs, Rogier Kievit, Anna McCarrey, Abdur Mustafa, Darren Price, David Samu, Jason R Taylor, Matthias Treder, Janna van Belle, Nitin Williams, Daniel Mitchell, Ethan Knights, Adam Attaheri, Dace Apsvalka, Maite Crespo-Garcia; Research Assistants: Lauren Bates, Tina Emery, Sharon Erzinçlioğlu, Andrew Gadie, Sofia Gerbase, Stanimira Georgieva, Claire Hanley, Beth Parkin, David Troy, Ina Demetriou, Will Duckett; Affiliated Personnel: Simon Fisher, Else Eising, Matt Bracher-Smith, Valentina Escott-Price, Tibor Auer, Lu Gao, Emma Green, Rafael Henriques; Research Interviewers: Jodie Allen, Gillian Amery, Liana Amunts, Anne Barcroft, Amanda Castle, Cheryl Dias, Jonathan Dowrick, Melissa Fair, Hayley Fisher, Anna Goulding, Adarsh Grewal, Geoff Hale, Andrew Hilton, Frances Johnson, Patricia Johnston, Thea Kavanagh-Williamson, Magdalena Kwasniewska, Alison McMinn, Kim Norman, Jessica Penrose, Fiona Roby, Diane Rowland, John Sargeant, Maggie Squire, Beth Stevens, Aldabra Stoddart, Cheryl Stone, Tracy Thompson, Ozlem Yazlik; and administrative staff: Dan Barnes, Marie Dixon, Jaya Hillman, Joanne Mitchell, Laura Villis.

## Data availability

Raw and minimally pre-processed MRI and behavioural data are available from https://camcan-archive.mrc-cbu.cam.ac.uk/dataaccess/. The raw data is publicly available for the purpose of scientific investigation subject to a data usage agreement. Analysis code is available at https://doi.org/10.6084/m9.figshare.34531299. All other materials supporting the findings of this study are available from the corresponding author upon request.

## Author contributions

D.J.M. and R.C. designed the project. D.J.M. performed analyses and wrote the first draft of the paper.

D.J.M. and R.C. revised the paper.

## Extended data

**Extended Data Figure 1.**
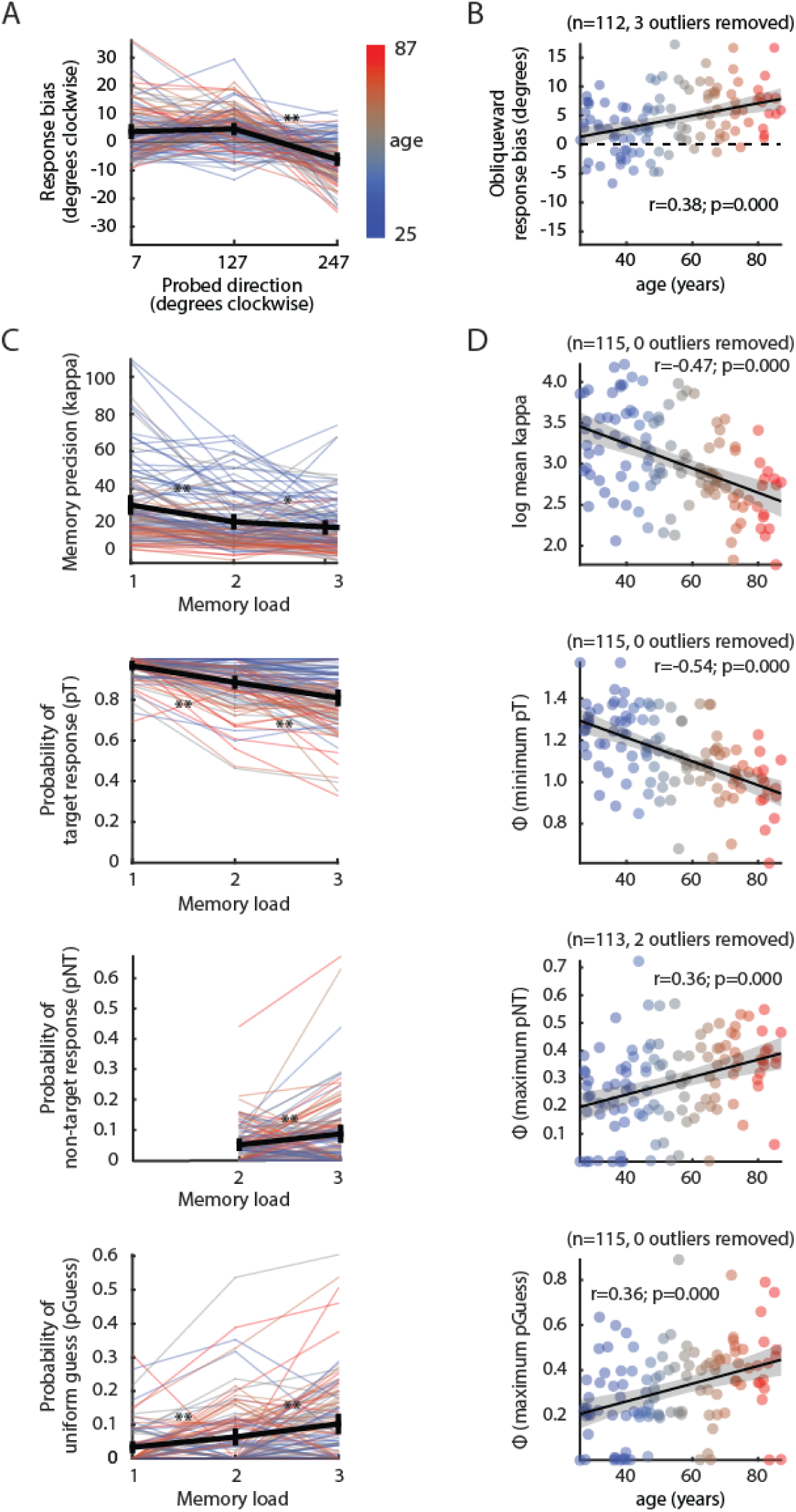
Mixture model decompositions of behavioural errors. (A) Preliminary model, assessing response bias as a function of motion direction (collapsed across memory load). (B) Summarising across motion directions, obliqueward bias increases with age. (C) Secondary model, per memory load, after subtracting obliqueward response bias. As memory load increases, the precision and likelihood of target responses decrease; the likelihood of non-target responses and random guesses increase. (D) Summarizing across memory loads, worse performance on each memory measure is associated with increasing age. In (A, C), thin lines represent individual participants, coloured by age; thick black lines represent the mean, with 95% confidence intervals. Asterisks mark significant differences between adjacent memory loads, * p<0.05, **p<0.01. In (B, D), black lines and grey error bands indicate linear trends with 95% confidence intervals. Φ indicates arcsine transform of proportional measures.

## Notes

### Competing Interest Statement

The authors have declared no competing interest.

